# Exactly solvable models of stochastic gene expression

**DOI:** 10.1101/2020.01.05.895359

**Authors:** Lucy Ham, David Schnoerr, Rowan D. Brackston, Michael P. H. Stumpf

**Affiliations:** School of BioSciences and School of Mathematics and Statistics, University of Melbourne, Parkville VIC 3010, Australia; Dept. Life Sciences, Imperial College London, SW7 2AZ, UK

## Abstract

Stochastic models are key to understanding the intricate dynamics of gene expression. But the simplest models which only account for e.g. active and inactive states of a gene fail to capture common observations in both prokaryotic and eukaryotic organisms. Here we consider multistate models of gene expression which generalise the canonical Telegraph process, and are capable of capturing the joint effects of e.g. transcription factors, heterochromatin state and DNA accessibility (or, in prokaryotes, Sigma-factor activity) on transcript abundance. We propose two approaches for solving classes of these generalised systems. The first approach offers a fresh perspective on a general class of multistate models, and allows us to “decompose” more complicated systems into simpler processes, each of which can be solved analytically. This enables us to obtain a solution of any model from this class. We further show that these models cannot have a heavy-tailed distribution in the absence of extrinsic noise. Next, we develop an approximation method based on a power series expansion of the stationary distribution for an even broader class of multistate models of gene transcription. The combination of analytical and computational solutions for these realistic gene expression models also holds the potential to design synthetic systems, and control the behaviour of naturally evolved gene expression systems, e.g. in guiding cell-fate decisions.

## I. INTRODUCTION

The need for models in cellular biology is ever-increasing. Technological and experimental advances have led not only to progress in our understanding of fundamental processes, but also to the provision of a great wealth of data. This presents enormous opportunities, but also requires further development of the mathematical and computational tools for the extraction and analysis of the underlying biological information.

The development of mathematical models for the complex dynamics of gene expression is of particular interest. The stochastic nature of the transcriptional process generates intracellular noise, and results in significant heterogeneity between cells; a phenomenon that has been widely observed in mRNA copy number distributions, even for otherwise identical cells subject to homogeneous conditions^1–6^. Modelling is key to understanding and predicting the dynamics of the process, and subsequently, to quantifying this observed variability. The Telegraph model, as introduced by Ko et al.^7^ in 1991, is the canonical model for gene expression and explains some of the variability observed in mRNA copy number distributions. This mathematical model treats gene promoter activity as a two-state system and has the advantage of a tractable stationary mRNA distribution^4,8,9^, enabling insights into limiting cases of the system^10^ and the behaviour of the distribution in general.

With the advancement of single-cell experiments, the deficiencies of the Telegraph model are increasingly apparent. The model does not account for more complex control mechanisms, nor more complex gene regulatory networks involving feedback. Recent work has shown that, while the Telegraph model is able to capture some degree of the variability in gene expression levels, it fails to explain the large variability and over-dispersion in the tails seen in many experimental datasets^11^. Furthermore, it has become evident that gene promoters can be in multiple states and may involve interactions between a number of regulatory factors, leading to a different mRNA synthesis rate for each state; such states may be associated with the presence of multiple copies of a gene in a genome, or with the key steps involved in chromatin remodelling^12–15^, DNA looping^16^ or supercoiling^17^. Mathematical models—and accompanying analytic or approximate solutions—that capture the kind of data arising from these more complex multistate dynamics are now essential.

Despite its deficiencies, the Telegraph model remains a useful framework around which more complicated models can be constructed and, as we will show here, provides a foundation from which to develop further analytical results. A number of extensions and modifications of the Telegraph model have emerged, the simplest of which is perhaps the extension to allow for a second basal rate of transcription—the “leaky” Telegraph model^11,18,19^. A further extension incorporating the additional effects of extrinsic noise is given in Refs. 11, 20, and 21. Several recent studies of gene expression have shown that the waiting time distribution in the inactive state for the gene promoter is non-exponential^22–24^, leading to the proposal of a number of “refractory” models^25–28^. A distinguishing feature of these models is that the gene promoter can transcribe only in the active state and, after each active period, has to progress through a number of inactive states before returning to the active state. An exact solution for the stationary distribution of a general refractory model is derived in Ref. 26, building upon the existing work of Refs. 25 and 27. In addition to direct extensions of the Telegraph model, many variants of the model have been proposed, some of which have been additionally solved analytically. Examples include a two-state model incorporating a gene regulatory feedback loop^29,30^; a three-stage model accounting for the discrepancy in abundances at the mRNA and protein level^31^; and more recently, a multi-scale Telegraph model incorporating the details of polymerase dynamics is solved in Ref. 32. Some approximation methods have been developed to solve more general multi-state models^19,33^.

Obtaining analytical solutions to models of gene transcription is a challenging problem, and there currently exist only a few classes of systems and a handful of special cases for which an analytic solution is known. A number of these have just been listed; see Ref. 34 for more information.

Existing methods for solving such models adopt a chemical master equation approach and typically employ a generating function method, in which the original master equation is transformed into a finite system of differential equations^35^. When an exact solution to the differential equations defining the generating function can be found, the analytical solution to the model can in principle be recovered by way of successive derivatives. A major complication in finding analytical solutions in this way is that typically even slight changes to the original model lead to considerably more complicated systems of differential equations. An additional challenge is that analytic solutions to models usually involve hypergeometric functions, which can lead to numerical difficulties in further computational and statistical analysis. Due to the lack of success in the search for analytic solutions, a significant amount of effort has been on stochastic simulation^36^ and approximation methods^34^. However, some of these approaches can be computationally expensive, making them unviable for larger systems, while most existing approximation methods are adhoc approximations that do not allow to control the approximation error. To address these obstacles, we present two alternative methods for solving certain classes of models of gene transcription.

The first approach offers a novel conceptualisation of a class of multistate models for gene transcription and allows us to extend a number of known solutions for relatively basic processes to more complicated systems. For a certain class of models, we are able to abstractly decompose the more complicated system as the independent sum of simpler processes, each of which is amenable to analytic solution. The distribution of the overall system can then be captured by way of a convolution of the simpler distributions.

The method applies to a wide range of models of stochastic gene expression. Examples include systems with multiple copies of a single gene in a genome, or systems with multiple promoters or enhancers for a single gene, each independent, and contributing additively to the transcription rate. Here we will provide analytical expressions for a multistate process consisting of a finite, exponential number of activity states, each with a particular mRNA transcription rate. Similar theoretical models have been employed to predictively model multistate processes^37,38^, however, the analyses in these are done numerically. Thus, our results enable exact solutions to some natural examples of multistate gene action, with an unlimited (though exponential) number of discrete states.

The second method draws as inspiration from the original derivation of the solution to the Telegraph model, given by Peccoud and Ycart^8^ in 1995, and provides an alternative approximative approach to solving master equations. The method is reminiscent of Taylor’s solution to differential equations^39^, using the differential equations to construct a recurrence relation for derivatives of the solution. We show how our approach can be used to recover the full solution to the previously mentioned multistate model to arbitrary numerical precision. However, the method is also applicable to more complex systems for which no analytic solutions exist in the literature. When an analytical solution is known, we are able to benchmark the computational speed of the two approaches. For some systems, the approximation method is in fact faster than the analytical solution, as the latter typically requires multiple numerical calculations of the confluent hypergeometric function.

This article is structured as follows. We begin in Section II by presenting a general class of multistate models that cover many examples of gene transcription models used in the analysis of experimental data. We explain how these multistate processes can be decomposed into simple processes acting independently, and how these can be solved analytically. We complete the section by giving some examples of previously-unsolved models. In Section III, we consider the effects of extrinsic noise, and show that the main results of Ref. 11 can be extended to these models. Next, in Section IV, we outline the strategy for turning differential equations into recurrence relations that can be computationally iterated to produce solutions of arbitrary accuracy. We then utilise the approach to solve some examples of gene transcription models for which no analytic solutions are currently known. Finally, we assess the accuracy and computational efficiency of our approach. Comparisons are made between our approximative approach, analytical solutions, and the stochastic simulation algorithm^40^.

## II. DECOMPOSING MULTISTATE PROCESSES

The standard method of solving models of gene transcription is to transform the chemical master equation into a system of ordinary differential equations by way of generating functions. We will observe that there are certain situations in which the need to solve these differential equations can be avoided, by decomposing into sums of independent simpler processes, each of which can be treated analytically. These can then be recombined to solve the original system.

We now briefly outline the content of this section. After first defining the leaky Telegraph model, we present a generalisation of this model that allows for a finite number of discrete activity states. As motivation, we will apply our decomposition method to the leaky Telegraph model, which has a known exact solution for the steady-state mRNA copy number distribution. We will demonstrate how efficiently and easily this solution can be derived using the new approach. A more formal exposition of the method will then be given. Novel results are presented in the remaining subsections, where we describe in detail how this approach can be used to solve more complex, previously unsolved systems.

### A. The leaky Telegraph model

The leaky Telegraph model has been considered in a number of previous studies such as Refs. 11, 18, 19, and 25, and incorporates the well known phenomenon of promotor leakage and basal gene expression^41–43^. In this model, the gene is assumed to transition between two states, active (*A*) and inactive (*I*), with rates *λ* and *µ*, respectively. Transcription of mRNA is modelled as a Poisson process, occurring at a basal background rate of *K*_*I*_ when inactive, and at a rate *K*_*A*_ > *K*_*I*_ when active. Degradation occurs as a first-order Poisson process with rate *δ*, and is assumed to be independent of the gene promoter state. A schematic of the system is given in Fig. 1. Observe that the standard Telegraph model corresponds to the special case for *K*_*I*_ = 0.

**FIG. 1.**
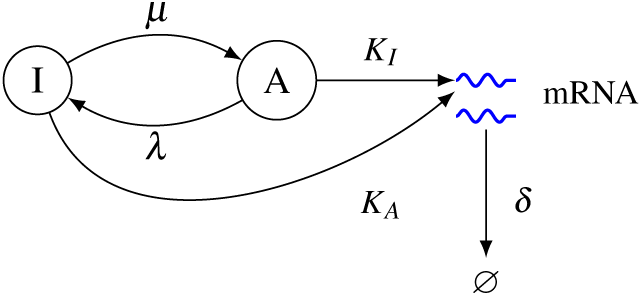
A schematic of the leaky Telegraph model. The nodes represent the state of the gene, labelled *I* and *A* for inactive and active, respectively. The mRNA are represented as blue wiggly lines. The parameters *λ* and *µ* are the rates at which the gene transitions between the two states *I* and *A*; the parameters *K*_*I*_ and *K*_*A*_ are the transcription rate parameters for *I* and *A*, respectively; the degradation rate is denoted by *δ*.

We recall here that the mRNA copy number from the leaky Telegraph process, as shown in Ref. 11, has the following steady-state distribution:

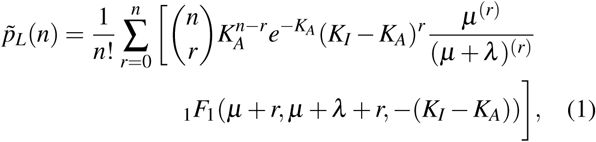

where 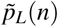 is the probability of observing *n* mRNA molecules in the system, _1_*F*_1_ is the confluent hypergeometric function^44^, and, for real number *x* and positive integer *n*, the notation *x*^(*n*)^ abbreviates the rising factorial of *x*. Note also that the rates are scaled so that *δ* = 1. When *K*_0_ = 0, we can recover the following well-known expression for the Telegraph process in the steady-state:

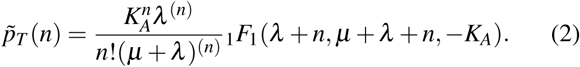

Throughout this work, we will refer to the probability mass functions 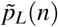 and 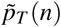 as the “leaky Telegraph distribution” and “Telegraph distribution”, respectively.

Another way to arrive at the same overall behaviour as the leaky Telegraph process is to consider a system consisting of two independent copies of the same gene, one lacking the control features of the other so that it transcribes constitutively at a constant rate of *K*_0_ := *K*_*I*_. The other copy of the gene is governed by a Telegraph process with transcription rate *K*_1_ := *K*_*A*_ − *K*_*I*_ when active (and 0 when inactive). Note that when the second gene is active, the combined transcription rate is *K*_0_ + *K*_1_ = *K*_*A*_, so this model is indistinguishable from the leaky Telegraph model; see Fig. 2 for a pictorial representation of this situation. The new interpretation is more easily modelled, as it is simply two independent systems, both of which are simpler than the original.

**FIG. 2.**
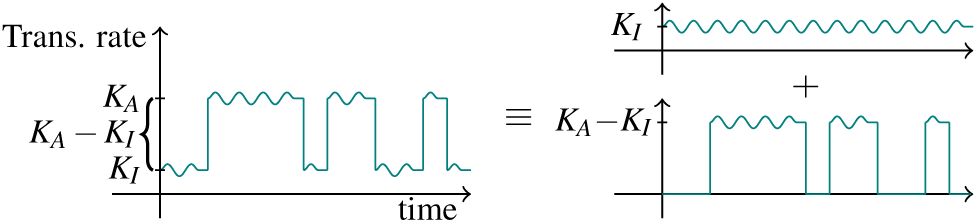
Decomposing the leaky Telegraph process (left) into the independent sum of a constitutive and Telegraph process (right).

We are able to make this observation more precise by way of analysis of the corresponding probability generating functions, defined as 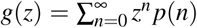, where *p*(*n*) is the stationary probability distribution. Let *P* be a random variable that counts the copy number of a constitutive process with constant rate *K*_0_, which is well-known to be Pois(*K*_0_) distributed at stationarity^45–47^, and let *T* be an independent random variable distributed by a Telegraph distribution with rates *λ,µ, K*_1_ = *K*_*A*_ − *K*_*I*_. Then the total copy number *X* for the binary system is simply the independent sum of *P* and *T*. A standard result in probability theory says that the probability generating function of a sum of two independent random variables is the product of the two component probability generating functions^48^ (Section 3.6(5)). The probability generating function for the random variable *P* is 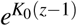 while the probability generating function for the Telegraph distributed *T* is _1_*F*_1_ (*λ,µ* + *λ, K*_1_(*z* − 1))^8^. So the probability generating function for the overall system is:

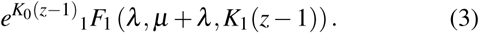

This solution, (3), should match the probability generating function yielding (1). From Ref. 11, the probability generating function for the leaky Telegraph model is given by:

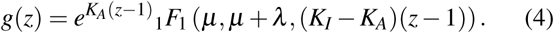

Multiplying (4) though by 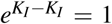, and applying Kummer’s transformation for the confluent hypergeometric function, we obtain (3), noting that *K*_0_ = *K*_*I*_ and *K*_1_ = *K*_*A*_ − *K*_*I*_.

### B. The 2^*m*^-multistate model

We now extend the leaky Telegraph process of the previous section to allow for a finite number of discrete activity states. For a non-negative integer *m*, consider a gene with *m* distinct activating enhancers *a*_0_, …, *a*_*m*−1_, where for each *i* ∈ {0,…, *m* − 1}, the enhancer *a*_*i*_, independently, is either bound or unbound to activators with rates *λ*_*i*_ and *µ*_*i*_ (resp.) and contributes, additively, with rate *k*_*i*_ to the overall transcription rate of the gene. So each *a*_*i*_ is either bound or unbound, leading to 2^*m*^ possible states for the activity of the gene, which we denote by 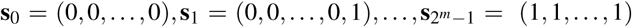, where for notational convenience **s**_*i*_ is the *m*-bit binary representation of the number *i* as a tuple. Tuples differing in just one position (that is, of Hamming distance *d*(**s**_*i*_, **s** _*j*_) = 1) will be said to be *adjacent*. Changes in state result from a binding or unbinding of an enhancer, from which it follows that state transitions will occur only between adjacent states.

We allow for a basal (leaky) rate of transcription *K*_*B*_, which is independent of the state of the enhancers. Then in each state **s**_*i*_, the mRNA are transcribed according to a Poisson process with a constant rate *K*_*i*_ = *K*_*B*_ + **k** · **s**_*i*_, for *i* ∈ {0, …, 2^*m*^ − 1}, where **k** = (*k*_*m*−1_, *k*_*m*−2_, …, *k*_0_) is the vector of mRNA transcription rates for the enhancers and · is the usual dot product. Note that for *i* = 0 (all enhancers unbound) this gives *K*_0_ = *K*_*B*_, while for *j* ∈ {0,…, *m* − 1}, we have 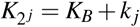.

The enhancers switch between bound and unbound with rates *λ*_*i*_ and *µ*_*i*_, respectively. This means that the system transitions between *adjacent* states 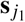 and 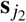 only: states differing in at most one bit, *i* say. This means that transition from 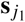 to 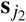 occurs stochastically at rate 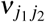 equal to either *λ*_*i*_ (if *j*_1_ < *j*_2_, meaning the *i*^th^ bit of *j*_1_ is 0) or *µ*_*i*_ (if *j*_2_ < *j*_1_). Degradation of mRNA occurs as a first-order Poisson process with rate *δ*, and is assumed to be independent of the promoter state. By a 2^*m*^*-multistate model*, we will mean a multistate model with 2^*m*^ states arising from the independent and additive transcription activity of *m* enhancers. We denote this model by **M**_*m*_. Figure 3 gives a depiction of the 2^*m*^-multistate process.

**FIG. 3.**
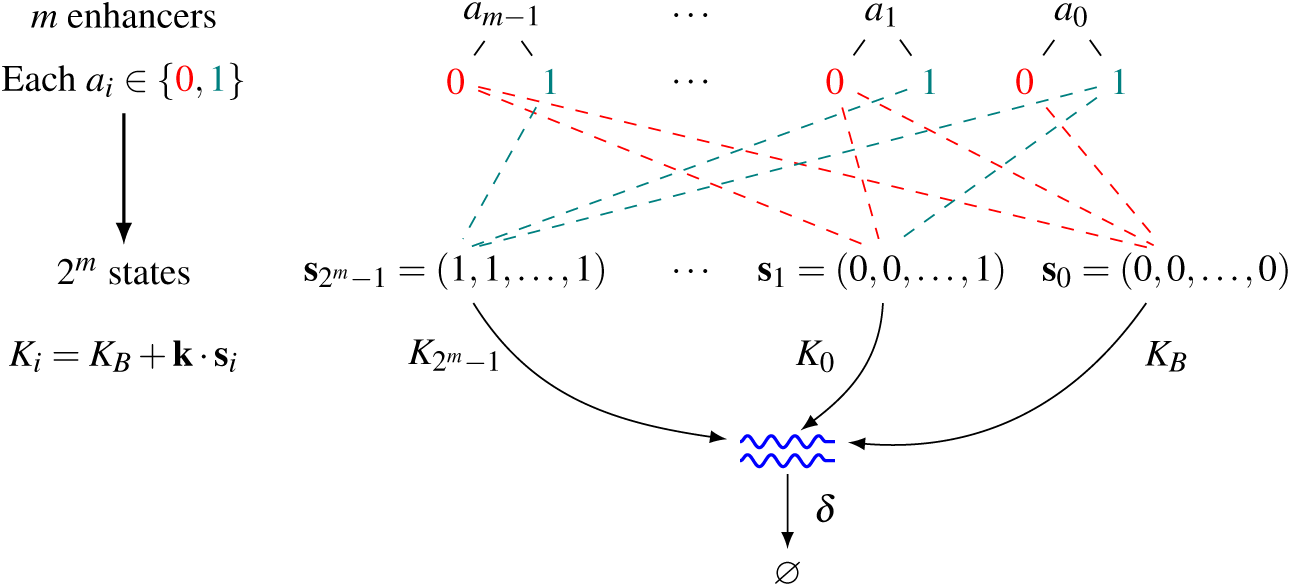
A diagrammatic representation of the 2^*m*^-multistate model, for *m* ∈ ℕ∪{0}. There are *m* enhancers, *a*_0_,…, *a*_*m*−1_, where each enhancer is either unbound (0) or bound (1): so *a*_*i*_ ∈ {0, 1}. This leads to 2^*m*^ possible activity states (*a*_*m*−1_, …, *a*_0_) for the gene; the state **s**_0_ = (0, 0, …, 0) corresponds to when all of the *a*_*i*_ are unbound, the state **s**_1_ = (0, 0, …, 1) corresponds to when *a*_0_ is bound, and all other *a*_*i*_ are unbound, while the state 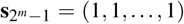 corresponds to when all of the *a*_*i*_ are bound. The gene transitions between these 2^*m*^ states, where switching events between adjacent states **s**_*i*_ and **s** _*j*_ occur at rate *ν*_*i j*_ (this is, however, not depicted in the above). Each state **s**_**i**_ has an associated synthesis rate, *K*_*i*_, equal to *K*_*B*_ + **k** · **s**_*i*_, where **k** = (*k*_*m*−1_, *k*_*m*−2_, …, *k*_0_) is the vector of mRNA transcription rates for the enhancers. Degradation occurs at rate *δ*, independently of the activity state.

If we let *M* represent the mRNA molecules, the reactions defining the 2^*m*^-multistate model can be summarised as:

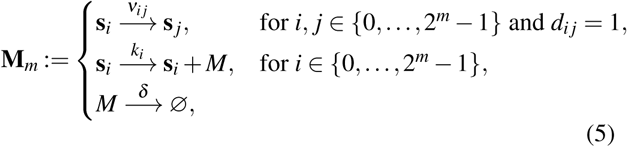

where *d*_*i j*_ is the Hamming distance between **s**_*i*_ and **s** _*j*_. Note also that *ν*_*i j*_, *k*_*i*_ ≥ 0, and *δ* > 0. The transition digraphs arising from the reactions defined in (5) can be visualised as hypercubes (*m*-dimensional cubes); see Fig. 4.

**FIG. 4.**
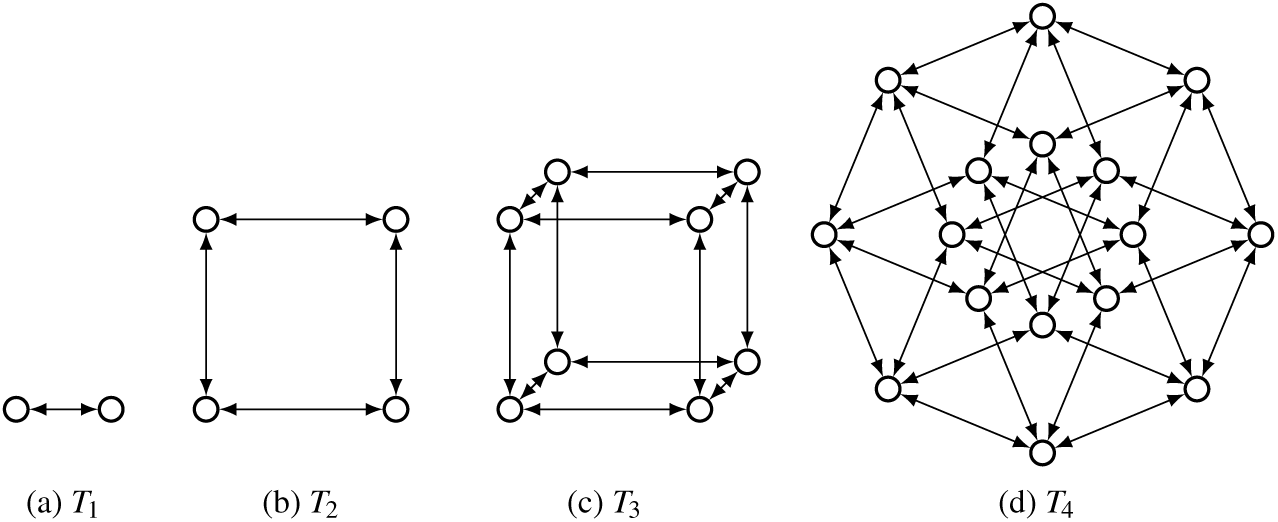
Transition digraphs arising from the 2^*m*^-multistate process. For *m* = 1, the transition digraph is a line segment, for *m* = 2 the transition digraph is a square, for *m* = 3 the digraph is a cube, and *m* = 4 corresponds to when the graph is a tesseract, and so on. Note that here we use *T*_*m*_ to denote the underlying transition digraph of the model **M**_*m*_.

Observe that the leaky Telegraph model coincides with the 2^1^-multistate model, **M**_1_, by setting *K*_*B*_ = *K*_*I*_ and *k*_0_ = *K*_*A*_ − *K*_*I*_. As a second example, the 2^2^-multistate model, **M**_2_, consists of two distinct activating enhancers *a*_0_ and *a*_1_, each with associated transcription rates *k*_0_ and *k*_1_, respectively. This leads to four possible activity states **s**_0_ = (0, 0), **s**_1_ = (0, 1), **s**_2_ = (1, 0) and **s**_3_ = (1, 1). The state **s**_0_ = (0, 0) corresponds to when both *a*_0_ and *a*_1_ are unbound. The state **s**_1_ = (0, 1) corresponds to when *a*_0_ is bound and *a*_1_ is unbound. Similarly, the state **s**_2_ = (1, 0) corresponds to when *a*_0_ is unbound and *a*_1_ is bound, and state **s**_3_ = (1, 1) corresponds to when both *a*_0_ and *a*_1_ are bound. The enhancers contribute additively to the overall transcription rate, so the rates for **s**_0_, **s**_1_, **s**_2_, **s**_3_ are *K*_0_ = *K*_*B*_, *K*_1_ = *K*_0_ + *k*_0_, *K*_2_ = *K*_0_ + *k*_1_, *K*_3_ = *K*_0_ + *k*_0_ + *k*_1_, respectively. Refer to Fig. 5(a) for a representation of the model **M**_2_.

**FIG. 5.**
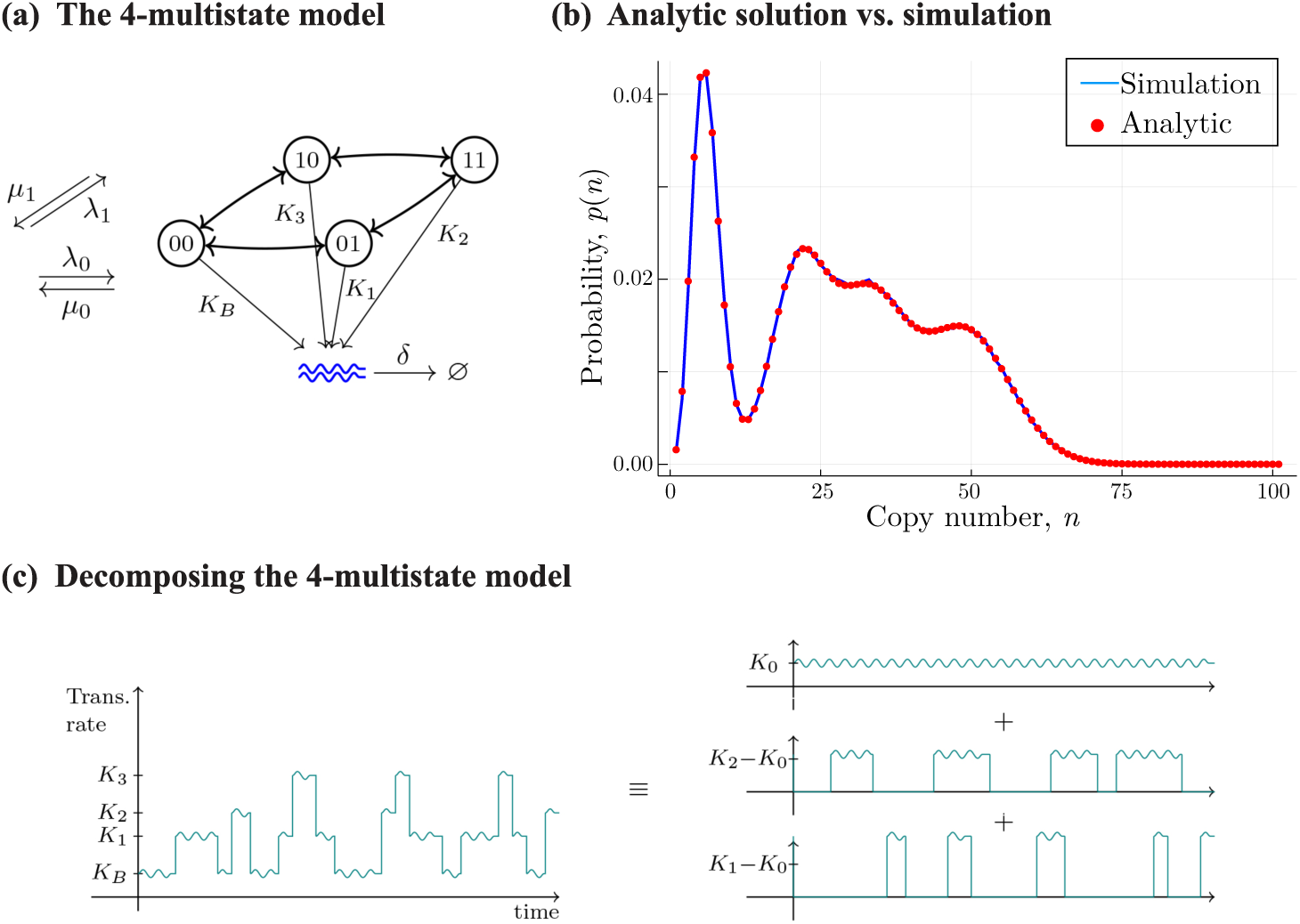
(a): A schematic of the 2^2^-multistate model (b): A comparison of the analytical solution for the steady-state mRNA distribution for the 2^2^-multistate model and the probability mass function from simulated data. The parameter values for both curves are *K*_0_ = 5, *k*_0_ = 21, *k*_1_ = 33 and *λ*_*i*_ = *µ*_*i*_ = 0.01, for *i* ∈ {0, 1}. The degradation rate *δ* is set to 1. (c): Decomposing the 2^2^-multistate model (left) into constitutive and Telegraph parts (right).

### C. Equivalent systems for M_*m*_

As with our motivating example, the 2^*m*^-multistate model has a courser-grained formulation obtained by disregarding the finer level information about the states of the activating enhancers. In this view, the 2^*m*^-multistate model can be equivalently thought of as a model for the transcriptional activity of

a. multiple enhancers of a single gene, where each enhancer has a particular additive effect on the mRNA transcription rate;
b. multiple independent copies of a single gene, where each copy of the gene is governed by a (possibly leaky) two-state system, and with possibly different transcription and switching rates;
c. a gene promoter with a finite, exponential number of states, where each promoter has an associated mRNA transcription rate, and transitions between states occur only between adjacent states.

The systems given in items (a), (b), and (c) have equivalent transition state diagrams, and therefore equivalent master equations. As an example, see Fig. 5(c) for a visualisation of the decomposition of the 2^2^-multistate model into simpler independent systems. It is easily seen that a random variable from the steady-state copy number distribution of the system given in (b) (which we know exists as the process has a finite transition digraph) is an independent sum of random variables from simpler distributions, namely Poisson and Telegraph distributions, each of which have exact solutions. It follows that a random variable from the steady-state mRNA copy number of the system given in (a) must also be amenable to the same decomposition, as we now show explicitly.

### D. Exact solution for the 2^*m*^-multistate model

Here we provide analytical solutions for any model of the class *ℳ* = {**M**_*m*_ | *m* ∈ ℕ_0_}. Let *m* ∈ ℕ_0_ and let *Y* be a random variable that counts the mRNA copy number of the 2^*m*^-multistate system in the steady-state. Recall that in this system the gene has *m* distinct activating enhancers *a*_0_, …, *a*_*m*−1_, each switching between the states bound or unbound, independently, with rates *λ*_*i*_ and *µ*_*i*_, and contributing additively with rate *k*_*i*_, for *i* ∈ {0, …, *m* − 1}, to the overall transcription rate of the gene. There is also a background rate of transcription *K*_0_ = *K*_*B*_. As before, we let *P* be a random variable for a constitutive process with constant rate *K*_0_ and, for each *i* ∈ {0, …, *m* − 1}, let *T*_*i*_ be a random variable distributed by a Telegraph distribution, 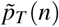, with rates *λ*_*i*_, *µ*_*i*_, *k*_*i*_. We have already seen that the leaky Telegraph system (which can be thought of as arising from a 2^1^-multistate process) has the associated random variable decomposition *X* = *P* + *T*, where *T* is Telegraph distributed with rate *k*_0_. It follows from the equivalence of (a) and (b) above that this can be generalised to:

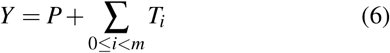

We can then directly write down the probability generating function for the 2^*m*^-multistate model as:

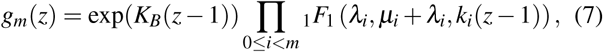

which is simply the product of a Poisson probability generating function and *m* Telegraph probability generating functions. The probability mass function is then recovered as 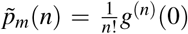, which by the extension of the general Leibniz rule to more than two factors gives:

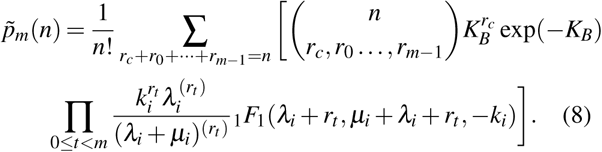

As an example of (8), the 2^2^-multistate model has probability mass function:

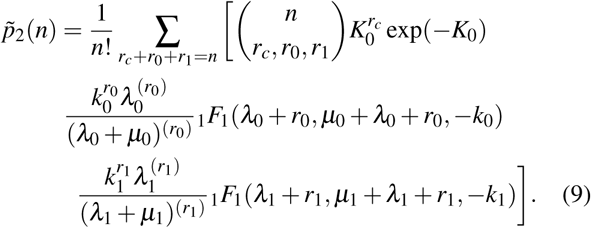

Figure 5(b) gives a comparison of 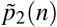 and the probability mass function from simulated data. A second example of (8) is a solution to the 2^3^-multistate model, depicted in Fig. 6. In both cases we observe excellent agreement of our method with stochastic simulations.

**FIG. 6.**
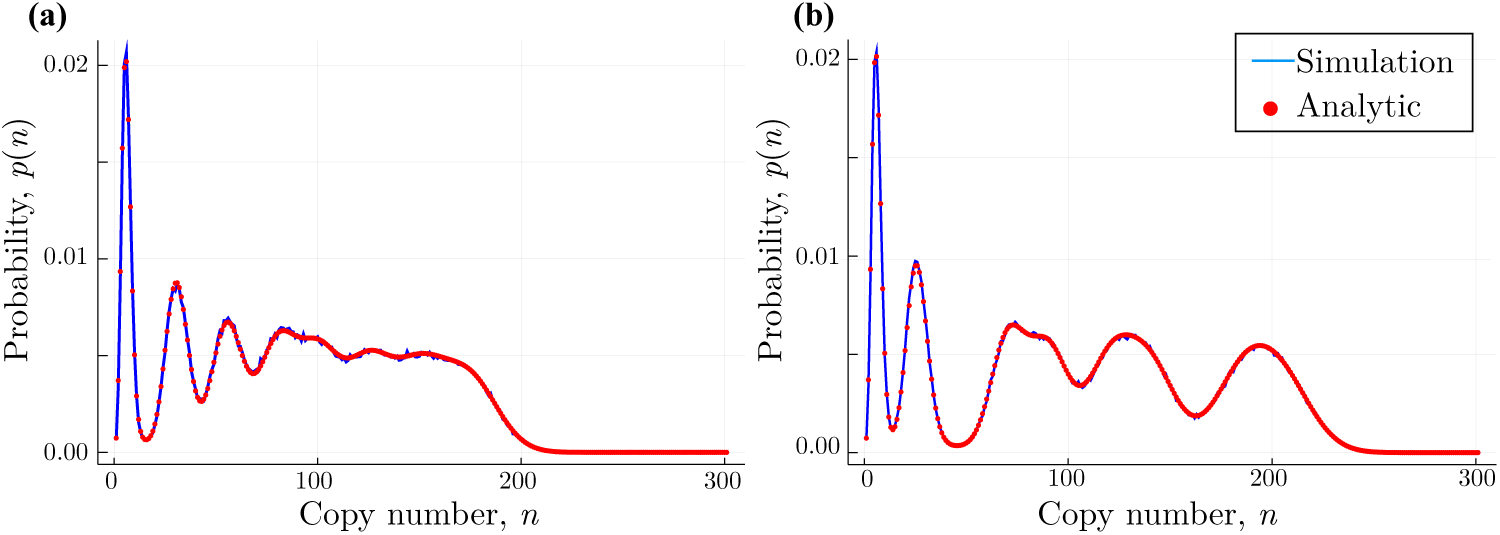
A comparison of the analytic solution to the 2^3^-multistate model and the probability mass function from simulated data. The parameter values for (a) are *K*_*B*_ = 5, *k*_0_ = 30, *k*_1_ = 50, *k*_2_ = 100 and *λ*_*i*_ = *µ*_*i*_ = 0.01, for *i* ∈ {0, 1, 2}. For (b) the parameters are *K*_*B*_ = 5, *k*_0_ = 20, *k*_1_ = 65, *k*_2_ = 115 and *λ*_*i*_ = *µ*_*i*_ = 0.01, for *i* ∈ {0, 1, 2}.

While the 2^1^-multistate model (corresponding to the leaky Telegraph model) has been solved before, to the best of our knowledge this is not the case for 2^*n*^-multistate models with *n* ≥ 2.

## III. EFFECTS OF EXTRINSIC NOISE

In a similar way to Ref. 11, we now jointly consider the effects of intrinsic and extrinsic noise on the probability distributions arising from the class of multistate models *ℳ* = {**M**_*m*_ | *m* ∈ ℕ_0_}. The results given in Ref. 11 are in the context of the leaky and standard Telegraph processes. We will show that the decomposition established in (6) allows us to straightforwardly extend the main analysis and results derived in Ref. 11 to *ℳ*. More specifically, we show that

a. intrinsic noise alone, as arising from the inherent stochasticity of the 2^*m*^-multistate process, never results in a heavy-tailed mRNA copy number distribution;
b. certain forms of extrinsic noise on at least one of the transcription rate parameters *k*_*i*_ (for *i* ∈ {0, …, *m* − 1}) or *K*_*B*_ is a sufficient condition for a heavy-tailed mRNA distribution;
c. the forms of this extrinsic noise are not limited to a specific distribution, but to any distribution that satisfies certain properties.

Our argument relies on moment generating functions, which for a random variable *X* with distribution *f*, is defined as M _*f*_ (*t*) := E(*e*^*tx*^) for *t* ∈ ℝ. We here take heavy-tailed to mean that the moment generating function is undefined for positive *t*. As in Ref. 11, the effects of extrinsic variability on the multistate systems is captured via a compound distribution; see Eq. 9 there. Items (a), (b), (c) follow directly from the following inequality for the moment generating function of the Telegraph distribution, 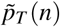: (Ref. 11, Eq. 16) for all positive *t*,

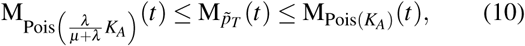

where M_*g*_ denotes the moment generating function for distribution *g*. We establish an analogous result for our multistate systems. First let *m* ∈ **N**_0_. Now since the moment generating function for **M**_*m*_ is simply the product of a Poisson moment generating function and *m* Telegraph moment generating functions it follows immediately from (10) that

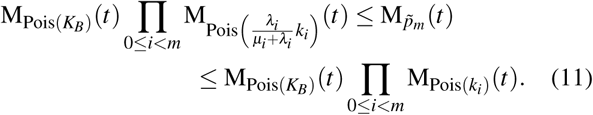

Now using the well-known result that the sum of *k* independent random variables *X*_*i*_ ∼ Pois(*γ*_*i*_) is a Pois 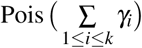 random variable, it follows that

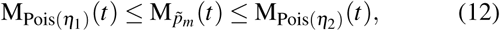

where 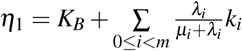 and 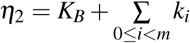. Now the arguments in Ref. 11 can be followed through identically: the moment generating function 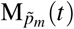 is bounded above by a Poissonian moment generating function that does not depend on any of the *λ*_*i*_ or *µ*_*i*_. Thus 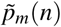 itself is not heavy tailed (showing (a)), and we require compounding of at least one of the *k*_*i*_ or *K*_*B*_ to make it so. Thus, (b) holds. On the other hand, *any* extrinsic noise on *K*_*B*_ or *k*_*i*_ that renders the moment generating function for Pois(*η*_1_) undefined, will also result in 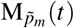 being undefined and the resulting compound distribution will be heavy tailed. A particular example is log normal noise on at least one of the *k*_*i*_ or *K*_*B*_.

## IV. THE RECURRENCE METHOD

We now introduce a recurrence method that can be used for approximating solutions to master equation models of gene transcription. We show that the method applies to a class of general multistate models in which any finite number of discrete states, along with arbitrary transitions between states, is allowed. In comparison to the 2^*m*^-multistate model, the number of states is no longer restricted to powers of two, and transitions between states are no longer restricted to adjacent states. In fact, the 2^*m*^-multistate model is a special case of this more general model. In addition, we show that the recurrence method is applicable to systems that are non-linear. We illustrate this by applying the method to the gene regulatory feedback model given in Ref 29.

We begin this section by introducing the master equation for the class of more general multistate models. Following this, we present a step-by-step breakdown of our recurrence method that can be used to provide a solution of desired accuracy to any multistate model from this class. We then illustrate the method through an example; see Subsection IV C. When an analytical solution can be obtained using the results of the previous section, we assess the computational efficiency and accuracy of the two solutions. We also make a comparison of our approach with the stochastic simulation algorithm. These results are given in Subsection IV F. Finally, we discuss the applicability and usefulness of our recurrence approach to other systems of gene transcription, including those involving feedback, as well as multiple stages, such as where protein copy number is modelled and the production of mRNA and protein are treated as separate stages.

### A. The *l*-switch model

In this section, we consider a system with *𝓁* distinct activity states **s**_1_, …, **s**_*l*_, where in each state **s** _*j*_, the mRNA are transcribed according to a Poisson process with a constant transcription rate *K*_*j*_. We assume switching events between states 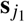 and 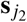 occur at rate 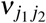 As before, the degradation of mRNA occurs as a first-order Poisson process with rate *δ*, and is assumed to be independent of the activity state. The reactions defining this *𝓁*-state model can be written as:

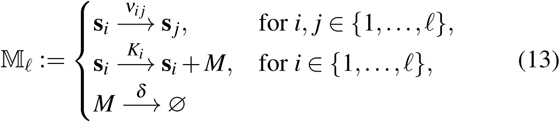

where again *M* represents the mRNA molecules, and *ν*_*i j*_, *k*_*i*_ ≥ 0, and *δ* > 0. We let 𝕄_*𝓁*_ denote the model defined by (13), and refer to 𝕄_*𝓁*_ as the *𝓁-switch model*.

Comparing with (5) we find that the 2^*m*^-multistate model agrees with the *𝓁*-switch model by setting *𝓁* = 2^*m*^, and setting *ν*_*i j*_ = 0 whenever *d*(**s**_*i*_, **s** _*j*_) > 1 and provided that the transcriptions rates adhere to the necessary additivity condition.

We are interested in analysing stationary distributions of the systems described in (13). Let *p*_*i*_(*n*) denote the probability at stationarity that the gene is in state *i* with *n* mRNA molecules, for *i* ∈ {1, …, *𝓁*}. The generalised chemical master equation for the probability mass function, *p*_*i*_(*n*), is given by:

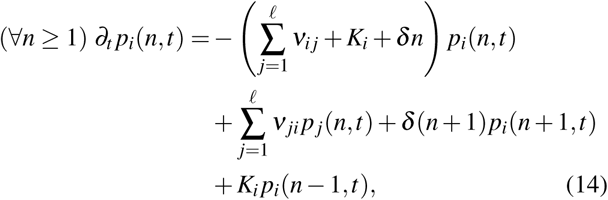

for *i* ∈ {1,…, *𝓁*}.

### B. The recurrence method

In the following, we give a step-by-step breakdown of the recurrence method applied to the system described in (14). After transforming the master equation (Step 1), the idea is to transform the resulting coupled system of differential equations into a closed system of recurrence relations in 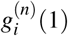 (Step 2), where *g*_*i*_(*z*) is the generating function of *p*_*i*_(*n*). Once the initial conditions *g*_*i*_(1) have been found, we can iteratively generate the 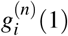 from the obtained recurrence relations, which in turn can be used to solve for *g*(*z*). This is detailed in Step 3. Finally, in Step 4, we can recover the solution for 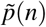 by way of 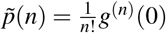.

A similar approach for transforming differential equations into recurrence relations is employed in Ref. 33 to find approximate solutions to *l*-switch models. It can be demonstrated that the final recurrence relations obtained from the transformation given here agree with those obtained in Ref. 33. While our method presents an alternative pathway to obtaining recurrence relations, the primary focus here will be on efficiency comparisons between our method, the previously obtained analytical results, and stochastic simulations, as well as applicability to non-linear models that do not belong to the *l*-switch class.

#### Step 1. Transform the master equation

As usual we can follow the generating function approach^35,49^ by defining, for each *i* ∈ {1, …, *𝓁*}, a stationary generating function:

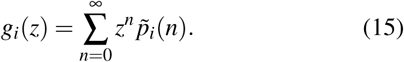

We transform the master equation (14) by multiplying through by *z*^*n*^ and summing over *n* from 0 to ∞, obtaining a system of *𝓁* coupled differential equations:

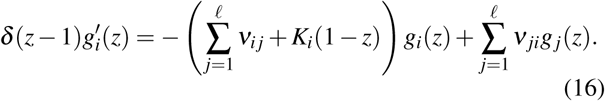

#### Step 2. Transform the differential equations into a system of first-order recurrence relations

Noting that in general the *k*^th^ derivative of a function of the form (*z* − 1)*h*(*z*) is (*z* − 1)*h*^(*k*)^(*z*) + *kh*^(*k*−1)^(*z*), by differentiating (*n* times) the equations in (16), we obtain for each *i* ∈ {1,…, *𝓁*}:

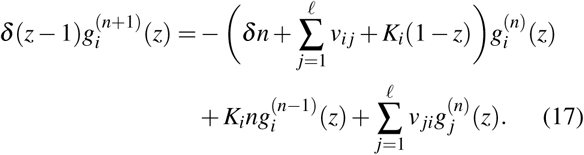

Evaluating each of the equations in (17) at *z* = 1, we immediately obtain the system of recurrence relations:

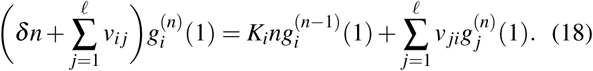

Rearranging each of these so that the LHS is equal to 0, the resulting system of equations can be represented in matrix form as:

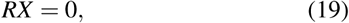

where *R* is a *𝓁* × 2*𝓁* matrix defined as:

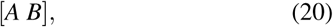

where *A* is an *𝓁* × *𝓁* matrix defined by:

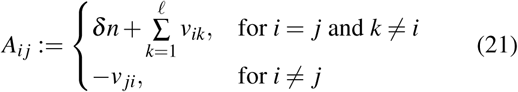

and *B* is a *𝓁* × *𝓁* matrix defined by:

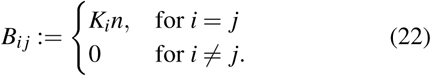

We also have

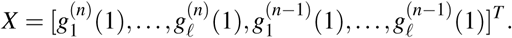

Eq. (18) constitutes a system of first-order linear recurrence relations and can be decoupled by applying Gaussian elimination to (19). This enables us to write each of the 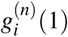 in terms of the 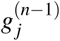. We let ℜ denote the recurrence relations obtained from the resulting matrix; as usual with Gaussian elimination, it is not practical to present a general form for the final simplified matrix.

#### Step 3. Iteratively solve the recurrence relations

We require first the initial conditions *g*_*i*_(1), for *i* ∈ {1, …, *𝓁*}. These can be found by evaluating equation (16) at *z* = 1, and solving the resulting system of linear equations. This system can be represented in matrix form as:

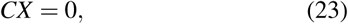

where the matrix *C* is defined as:

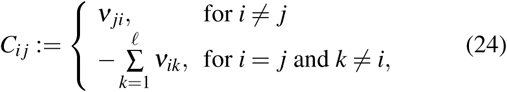

and we have

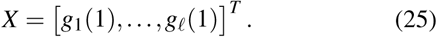

Now, as 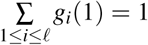, the *g*_*i*_(1) are found as a normalised version of Null(*A*), which is necessarily one dimensional for these multistate systems.

Now, if *g*_*i*_(*z*) is considered as a power series in (*z* − 1), then the coefficient of (*z* − 1)^*n*^ is 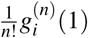. Thus, the coefficients 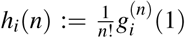 may be generated by iteration using the first-order recurrence relations, ℜ, obtained from (20). One way to do this would be to simply solve the recurrence relations for *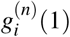* and to subsequently divide by *n*!. However, this involves computing a large number 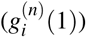 and dividing it by another large number (*n*!), which can lead to numerical problems. Instead, we transform the recurrence relations for 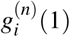 into recurrence relations for the *h*_*i*_(*n*).

Computationally, we may store the values for *h*_*i*_(*n*) in a system of *𝓁* lists from which we can compute the coefficients *h*(*n*) = ∑_*i*_ *h*_*i*_(*n*) of (*z* − 1)^*n*^ in the expansion of *g*(*z*) as a power series in (*z* − 1). So,

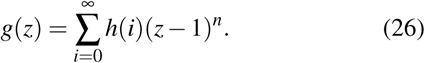

#### Step 4. Recover the stationary probability distribution

As *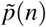* is the coefficient of *z*^*n*^ in the expansion of *g*(*z*) as a power series in *z*, we are able to recover 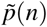 by way of 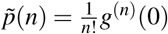. It then follows from (26) that

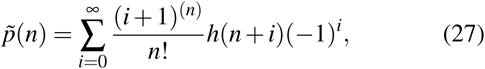

where again *x*^(*n*)^ denotes the rising factorial. For practical implementations, we truncate the sum in (27) after a certain number of terms. The distribution 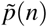 typically becomes negligibly small for *n* larger than a certain value, say 100. Computing *h*(*n*) up to *n* = 500 for example would thus leave at least 400 terms for approximating 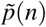 for *n* 100. For the studied example systems we found that 100−350 terms were sufficient to achieve accurate results.

### C. Example 1: the 3-switch model

We illustrate the recurrence method for the case *l* = 3.

*Steps 1 and 2*. From (20), the matrix, *R* := [*A B*] is comprised of:

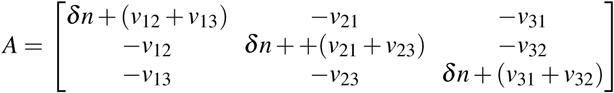

and

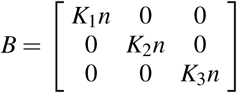

We also have that

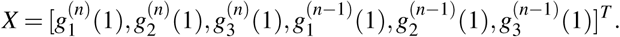

After applying Gaussian elimination to *R*, we obtain the following system of first-order linear recurrence relations, for all *i* ∈ {1, 2, 3} and *j < k* from {1, 2, 3}\{*i*}:

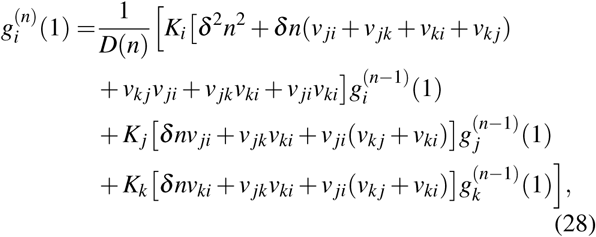

where

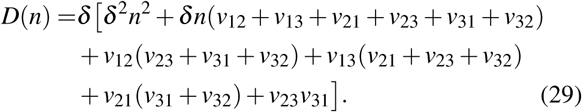

*Step 3*. We first find the initial conditions. From (24), the matrix, *C*, is given by:

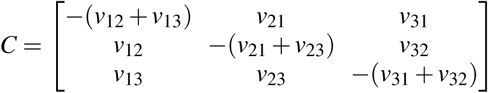

It can be shown that the null space, Null(*C*), is given by:

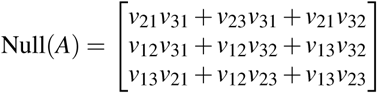

Thus, the initial conditions are:

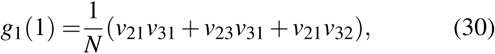

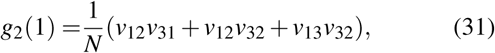

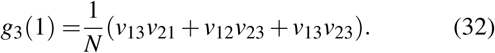

Here *N* is the normalisation constant and is equal to 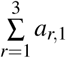 where each *a*_*r*,1_ is an element of the matrix Null(*A*).

Next, we need to compute the coefficients 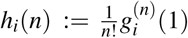 of *g*_*i*_(*z*) considered as a power series in (*z* − 1). As mentioned in Step 3 above, we transform the recurrence relations for 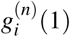 into recurrence relations for the *h*_*i*_(*n*). These can be obtained from (28) by dividing the RHS of each equation by *n*: for each {*i, j, k*} = {1, 2, 3},

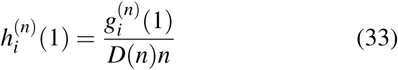

*Step 4*. Finally we recover the stationary distribution, 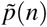. For a given set of parameter values, we generated a list of values for *h*(*n*) of length 500, and used this to approximate the probability mass function (27) from approximately 400 terms. The results for two different sets of parameter combinations are given in Fig. 7.

**FIG. 7.**
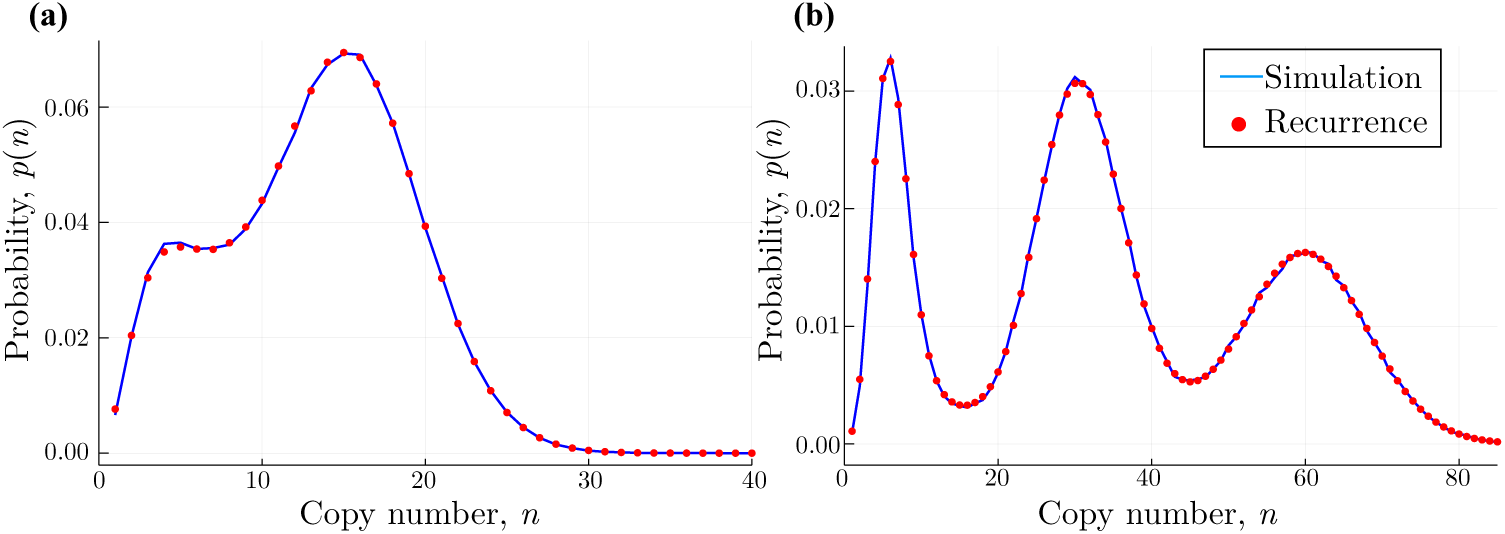
Stationary distributions of the 3-switch model in Section IV C. The figure shows the results obtained from stochastic simulations and from the recurrence method describes in Section IV. Figure (a) has parameters *v*_12_ = *v*_21_ = *v*_31_ = 0.1, *v*_13_ = 0.4, *v*_32_ = 0.9, and *K*_1_ = 2, *K*_2_ = 16, *K*_3_ = 5. Figure (b) displays a trimodal distribution with parameters *v*_12_ = *v*_13_ = 0.045, *v*_21_ = *v*_32_ = 0.035, *v*_31_ = *v*_23_ = 0.015, *K*_1_ = 5, *K*_2_ = 30, *K*_3_ = 60. We used 300 (a) and 305 (b) terms respectively for the recurrence method to approximate the sum in (27).

### D. Discussion of wider applicability

So far, we have seen how the recurrence method can be applied to linear *l*-switch models. In this section, we discuss the wider applicability of the method, and demonstrate that it can in principle also be applied to certain non-linear systems. As an example, we show that the method can be used to approximate the solution of the gene regulatory feedback model given in Ref. 29.

We would like to point out that the applicability to this system is limited, however. Application of the recurrence method to the feedback model relies on the initial conditions being supplied, and these are currently derived from the known analytic solution. This apparent circularity does not undermine the overall approach. The problem of solving chemical master equation systems can essentially be broken into two separate tasks: solving for the initial conditions for the generating function and solving for the probability mass function. Indeed, there are situations where the initial conditions of a given system can be easily found, but the probability mass function remains unknown; the *𝓁*-model is an example. On the other hand, there are situations where a general form for the probability mass function can be stated for a given set of initial conditions, but the initial conditions for the system are not yet known; this is the situation presented in Ref. 29. For the feedback model considered here, we are able to illustrate applicability to one half of the problem: for a given set of initial conditions for the generating function we can provide an approximation to the probability mass function.

We now apply the recurrence approximation to the system of differential equations arising from the feedback loop in Ref. 29. In this model of a single gene, the protein produced can bind to the promoter of this same gene, regulating its own expression. The process is modelled by recording only the number of proteins and the state of the gene promoter, which can be either bound or unbound. Here the rate of production of proteins depends on the state of the promoter region of the gene. Following Ref. 29, we let *r*_*u*_ and *r*_*b*_ denote the protein production rates for the bound and unbound states, respectively. We let *k* _*f*_ be the degradation rate of proteins, and let *k*_*b*_ be the degradation rate of the bound protein. Also letting *D*_*u*_, *D*_*b*_ denote the unbound and bound states of the gene promoter, respectively, and *P* denote the protein, the reaction scheme for the process can be written as:

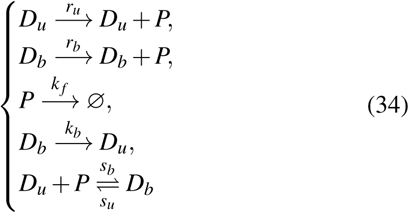

We refer the reader to Ref. 29 for further details of the model including the associated master equation (Ref. 29, Eq. 8). Applying the recurrence method to the feedback model (34), we obtain an approximation to the probability mass function. For two sets of parameter values, we compare this with the known analytic solution and present the results in Fig. 8. For details of the application of the method, including the derivation of the recurrence relations used for the approximation see Section A 2 of the appendix. The analytic solution for this model is given as equation (A23) in the appendix.

**FIG. 8.**
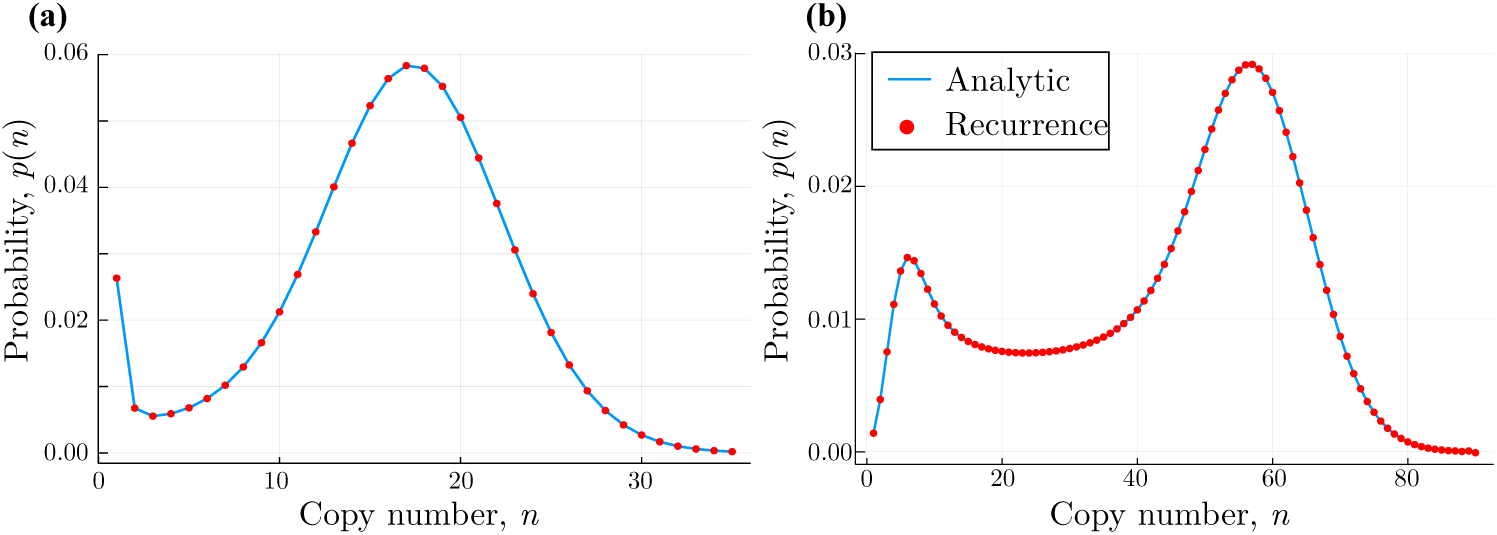
Stationary distributions of the feedback model in Section IV D. We compare the analytic solution with the results obtained from the recurrence approximation. Figure (a) has parameters *ρ*_*u*_ = 0.125, *ρ*_*b*_ = 20, *θ* = 0.5, *σ*_*u*_ = 0.2, *σ*_*b*_ = 0.35, and (b) has parameters *ρ*_*u*_ = 60, *ρ*_*b*_ = 6, *θ* = 0, *σ*_*u*_ = 0.5, *σ*_*b*_ = 0.004. We used 110 (a) and 300 (b) terms, respectively, for the recurrence method to approximate the probability mass function.

To this point, our discussion of the applicability of the recurrence method has concerned only chemical master equation models with a two-dimensional state space (tuples consisting of the state of the gene and the mRNA copy number or protein). Preliminary investigations suggest the method can be extended to models of gene transcription giving rise to higher dimensional state spaces. The three-stage model of Shahrezaei and Swain^31^ is an example. In this model, the production of mRNA and protein are treated as separate processes, resulting in a three-dimensional state space. In such cases, the recurrence method gives rise to equations containing partial derivatives. This makes the task of finding the initial conditions for the recurrence approach even more challenging. A characterisation of when the recurrence method is applicable to chemical master equation models will be the focus of future work.

### E. Notes on numerical implementation

We found that the recurrence method can become numerically unstable when solved with standard numerical precision for some of the studied example systems and parameter values. However, for sufficiently larger numerical precision we are able to obtain numerically stable and accurate results for all systems. We used Mathematica for the numerically difficult cases and provide example code in the supplementary information.

### F. Assessment of computational efficiency

We now assess the computational efficiency of the recurrence method by examining how the computational time scales with system size. We do this for the leaky Telegraph model, using the analytical solution to verify the accuracy of the result. As displayed by the example in Fig. 9(a), we find the two methods to be in complete agreement provided that a sufficient number of terms are considered in the expansion. For all of the models considered in this paper, the maximum copy number that must be considered *n*_max_, is directly related to the largest rate, in this case *K*_1_. We therefore apply the method for a number of *K*_1_, some examples of which are displayed in Fig. 9(b). We find that for each case a different number of terms in the expansion, and consequently a different computational cost, are required (Figs. 9(c), (d)). Scaling of the number of terms is observed to be approximately linear while scaling of the run-time is polynomial of 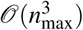.

**FIG. 9.**
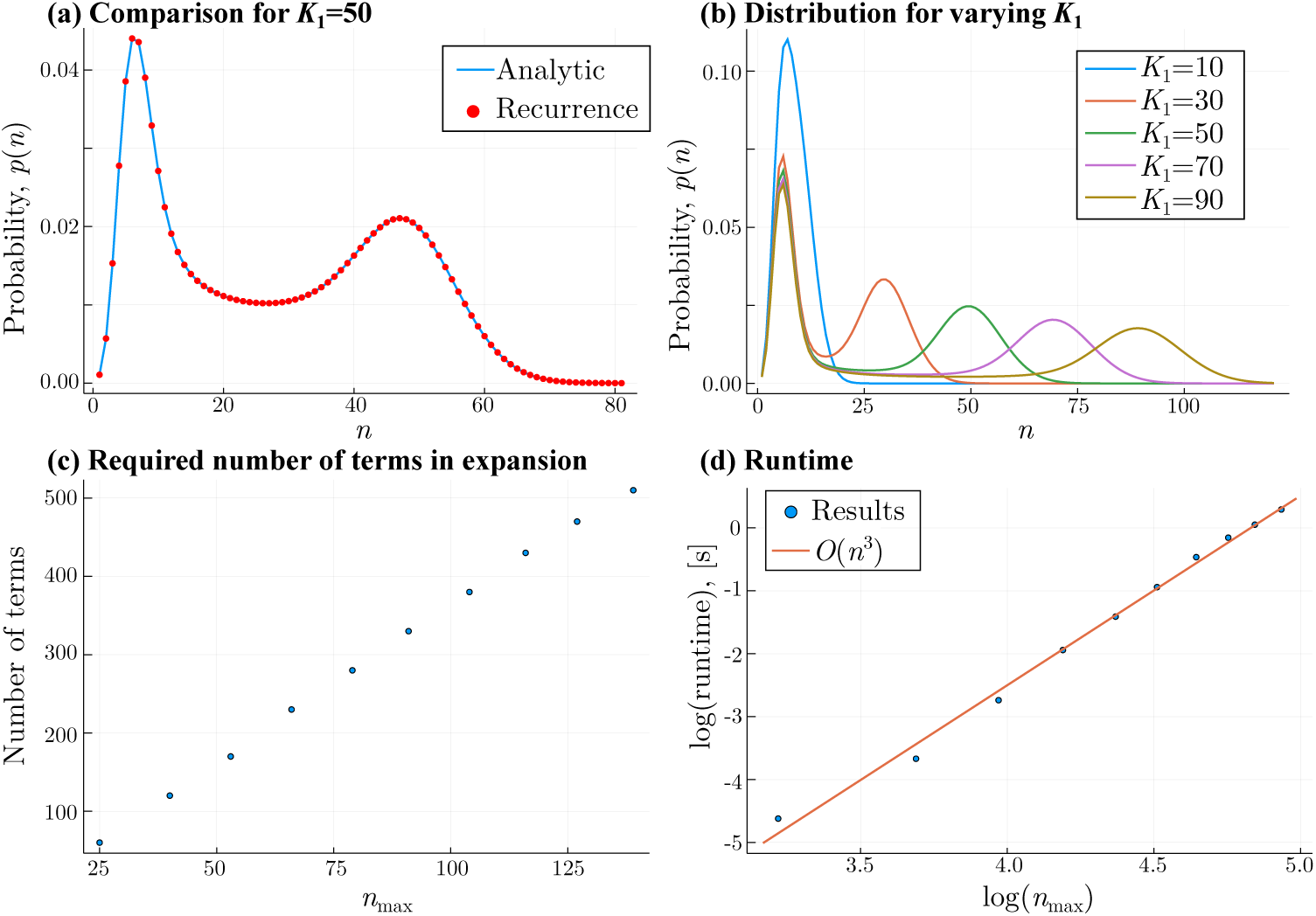
Scaling of the computational cost with increasing system size for the two-state model. In Fig. (a), we compare the analytic probability mass function for the leaky Telegraph model with the approximation obtained by recurrence method. The parameters are *λ* = 0.1, *µ* = 0.1, *K*_0_ = 5 and *K*_1_ = 50 and we used 270 terms to approximate the sum in (27) for the recurrence method. In (b), we plot the different leaky Telegraph distributions obtained by varying the parameter *K*_1_. Figure (c) shows the number of terms required in the recurrence expansion as a function of the maximum copy number with non-negligible probability in the system. We observe a linear relationship. Figure (d) shows that the runtime of the recurrence method scales roughly as the cube of the maximum copy number.

Interestingly, we observe in the numerical examples that the main computational cost of the recurrence method does not occur from solving the recurrence relation itself, but instead from the subsequent reconstruction of the probability mass function according to (27). This suggests that the computational cost of our method should be similar to or less than the computational cost when the analytic solution to the generating functions is known. For the feedback model studied above we observe that this is indeed the case.

For systems where no analytic solution is known one typically utilises the SSA to simulate sample trajectories from the stochastic process. Use of the SSA to obtain distributions is notoriously expensive Ref.34, and requires a sufficiently large number of statistically independent samples to be obtained.

For the results shown in Fig. 7 for the 3-switch model, for example, we found that simulating 300, 000 samples using our implementation of the SSA was about two orders of magnitudes slower than the recurrence method (∼ 57 (a) and ∼ 155 (b) seconds using the SSA, respectively, versus ∼ 0.1 (a) and ∼ 2 (b) seconds using the recurrence method.).

It is worth noting that even for systems for which analytic solutions exist, the solutions are typically expressed in terms of so-called “special functions” such as Bessel or Hypergeometric functions, which still need to be evaluated numerically and often by recurrence relations or series expansions Ref. 50. As an example see the probability mass function of the feedback model (A23) which involves a sum over confluent Hypergeometric functions. Such solutions can hence still be numerically expensive if one needs to evaluate over many points in state space. Indeed, we find that our recurrence method is considerably faster for approximating the probability mass function of the feedback model than evaluating the analytic expression.

## V. CONCLUSION

Despite the importance of stochastic gene expression models in the analysis of intracellular behaviour, their mathematical analysis still poses considerable challenges. Analytic solutions are only known for special cases, while few systematic approximation methods exist and stochastic simulations are computationally expensive. In this work, we provide two partial solutions to these challenges.

In this first part, we developed an analytic solution method for chemical master equations of a certain class of multistate gene expression models, which we call 2^*m*^-multistate models. The method is based on a decomposition of a process into independent sub-processes. Convolution of the solutions of the latter, which are easily obtained, gives rise to the solution of the full model. The solutions can be derived straightforwardly and are computationally efficient to evaluate. While most existing solutions only apply to specific models, our decomposition approach applies to the entire class of 2^*m*^-multistate models. This class covers a wide range of natural examples of multistate gene systems, including the leaky Telegraph model.

In the second part of this work, we derived a recurrence method for directly approximating steady-state solutions to chemical master equation models. The method was employed to approximate solutions to a broad class of linear multistate gene expression models, which we term *𝓁*-switch models, most of which currently have no known analytic solution. In particular, the class of *𝓁*-switch models to which the recurrence method applies is more general than (and contains) the class of 2^*m*^-multistate models to which the analytic method applies. The recurrence method is not limited to linear *𝓁*-switch models, however, but can also be applied to certain non-linear systems. Specifically, we have shown that given some known initial conditions, the method can be used to approximate the solution to a gene regulatory network with a feedback loop. For all studied systems, we found an excellent agreement of the recurrence method and analytic solutions or stochastic simulations.

In cases where no analytic solution is known, the recurrence method was found to significantly outperform stochastic simulations in terms of computational efficiency. Even in cases where an analytic solution exists, we found the method to be computationally more efficient than the evaluation of the analytic solution. While a characterisation of precisely which models the method is applicable to is not yet known, the results presented here suggest the recurrence method is an accurate and flexible tool for analysing stochastic models of gene expression. We believe that together with the presented analytical method it will contribute to a deeper understanding of the underlying genetic processes in biological systems.

## Supporting information

Sample Mathematica code

## SUPPLEMENTARY MATERIAL

See the supplementary material for sample Mathematica code for each application of the recurrence method considered in the present paper.

## ACKNOWLEDGMENTS

LH and MPHS gratefully acknowledge support from the University of Melbourne DVCR fund; RDB and MPHS were funded by the BBSRC through Grant BB/N003608/1; DS was funded by the BBSRC through Grant BB/P028306/1.

## Appendix A

### 1. Recurrence relations for the leaky Telegraph model

The following recurrence relations and initial conditions, obtained by applying the recurrence method to the leaky Telegraph model, can be used to give an approximation to the probability mass function (1).

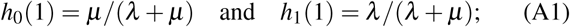

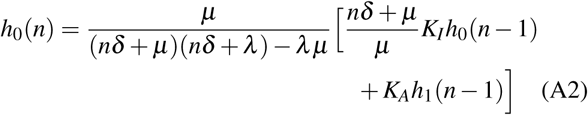

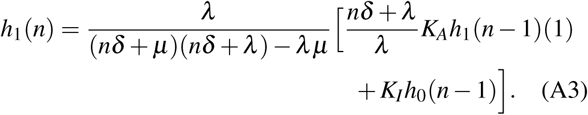

### 2. Recurrence relations for the feedback model

We provide details of the recurrence approximation applied to the system of differential equations arising from the feedback loop in Ref. 29. For convenience we list here the dimensionless variables that appear there:

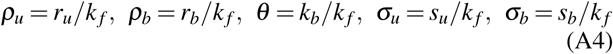

and the parameters:

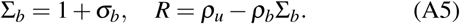

*Step 1.* The conveniently transformed equations from Ref. 29 (Equations (12) and (13)) are:

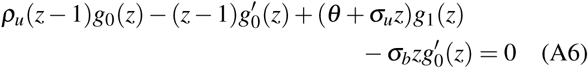

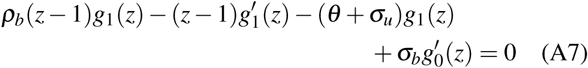

*Step 2.* Differentiating (*n* times) the equations (A6) and (A7) gives:

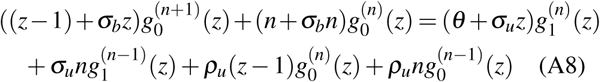

and

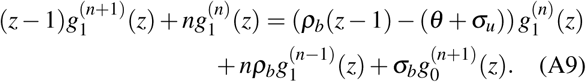

Evaluated at *z* = 1, we immediately obtain the recurrence relations

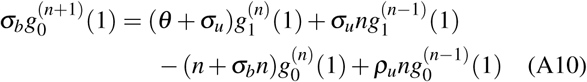

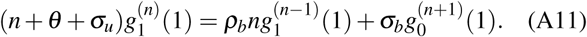

Substituting (A10) into (A11) leads to the following system of second-order linear recurrence relations: for *n* ≥ 1,

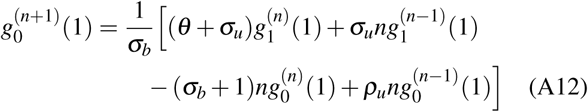

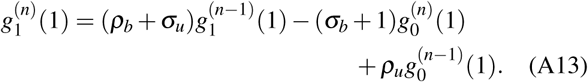

*Step 3.* Provided that the initial conditions *g*_0_(1), *g*_1_(1) and 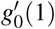 are known (here 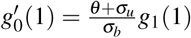), the values of 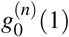 and 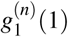 can be obtained by iteration. In this example, we use the initial conditions calculated from the known analytic solution (see Section A 3 for details of the initial conditions and the analytic solution).

*Step 4.* As before, we use a simplification that avoids the computational issue of calculating one huge number 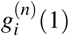, and then dividing by another huge number *n*!. The actual coefficients 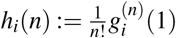 may be generated by iteration using the following recurrence relations obtained from (A12) and (A13):

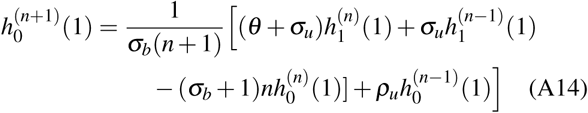

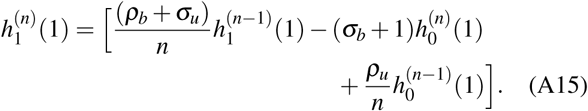

Note that here Equation (A14) is obtained from (A12) by dividing each term *g*_*i*_(*n* + 1 − *j*) by (*n* + 1)_(*j*+1)_, and similarly, Equation (A15) is obtained from (A13) by dividing each term *g*_*i*_(*n – j*) by *n*_(*j*)_, where *x*_(*i*)_ denotes the falling factorial of *x*.

*Step 5.* We are now able to recover the stationary probability mass function. Again, we generate a list of values for *h*(*n*) and use this to approximate the probability mass function (27). A comparison of the approximate solution and the known analytic solution is given in Figure 8.

### 3. Explicit solution to the feedback model

Here we provide an explicit solution for the stationary probability mass function of the regulatory feedback loop considered in Section IV D. From Refs. 29 and 30, the exact solutions for the generating functions are given by:

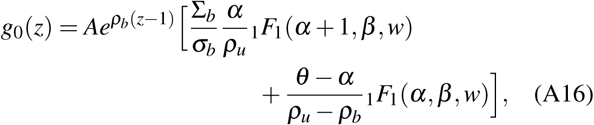

and

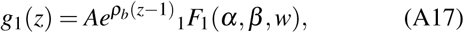

where

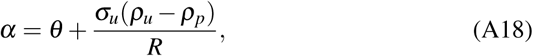

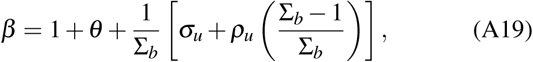

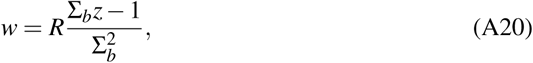

and

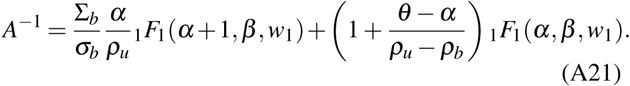

Here *w*_1_ denotes *w* evaluated at *z* = 1. The generating function is then obtained by *g*(*z*) = *g*_0_(*z*) + *g*_1_(*z*). So,

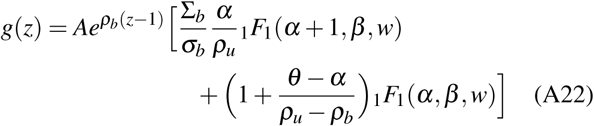

It follows that the stationary probability mass function is:

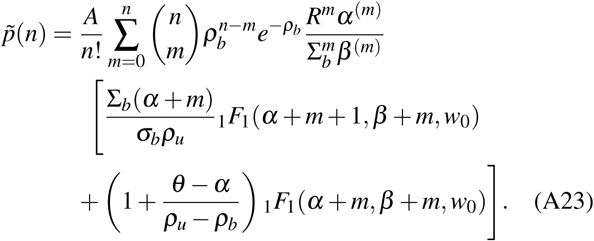

Again from (A22), the initial conditions *g*_0_(1) and *g*_1_(1) are:

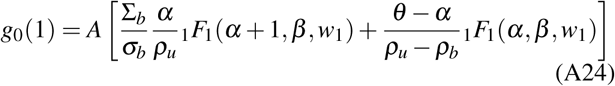

and

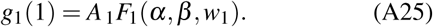

