## Supplementary material for "Exactly solvable models of stochastic gene expression": Sample Mathematica code

In[•]:=
```
Clear["Global`*"]
```

In[•]:=
```
SetDirectory@NotebookDirectory[];
```

### Two-state model

#### Definitions

##### Equations for recurrence method

```
(*Implement recurrence relation given in Equations (A1)-(A3)*)
```

$$\text{recurrence} := \Big\{ h0[n] == \frac{\mu}{(n\,\delta + \mu)\ (n\,\delta + \lambda) - \lambda\,\mu} \left( \frac{n\,\delta + \mu}{\mu}\ K0\ h0[n-1] + K1\ h1[n-1] \right),$$
$$h1[n] == \frac{\lambda}{(n\,\delta + \mu)\ (n\,\delta + \lambda) - \lambda\,\mu} \left( \frac{n\,\delta + \lambda}{\lambda}\ K1\ h1[n-1] + K0\ h0[n-1] \right),$$
$$h0[0] == \mu / (\lambda + \mu),$$
$$h1[0] == \lambda / (\lambda + \mu) \Big\}$$

In[•]:=
```
(*construct pmf from h list according to Equation (27) *)
```

$$p[h\_, n\_] := \frac{1}{n!}\ \text{Sum}\left[\text{Pochhammer}[i, n]\ h[[n+i]]\ (-1)^{i+1}, \{i, 1, \text{Length}[h] - n - 1\}\right]$$

##### Analytic solution

In[•]:=

$$\text{analytic}[n\_] := \frac{1}{n!}\ \text{Sum}\Big[\text{Binomial}[n, r]\ K1^{n-r}\ \text{Exp}[-K1]\ (K0 - K1)^{r}$$
$$\frac{\text{Pochhammer}[\mu, r]}{\text{Pochhammer}[\lambda + \mu, r]}\ \text{Hypergeometric1F1}[\mu + r, \lambda + \mu + r, -(K0 - K1)], \{r, 0, n\}\Big]$$

#### Example - Plot Figure 9 (a)

In[•]:=
```
(*set parameter values*)
δ = 1;
μ = 1/10;
λ = 1/10;
K0 = 5;
K1 = 50;
```

In[•]:=
```
(*perform recurrence relation*)
nest = RecurrenceTable[recurrence, {h0, h1}, {n, 0, 1000}];

(*construct pmf*)
hList = nest[[ ;; , 1]] + nest[[ ;; , 2]];
tay = {#, p[hList, #]} & /@ Join[Range[0, 10], Range[13, 80, 5]];
```

In[•]:=
```
(*compute analytic solution*)
ana = {#, analytic[#]} & /@ Range[0, 80];
```

In[•]:=
```
(*plot results*)
Show[ListLinePlot[ana, PlotRange → All, PlotStyle → Thickness[0.008],
   Frame → True, PlotLegends → {"analytic solution"},
   FrameLabel → {Style["n", 15], Style["p(n)", 15]}],
  ListPlot[tay, PlotStyle → {Red, PointSize → 0.02}, PlotRange → All,
   PlotLegends → PointLegend[{Red}, {"recurrence method"}]]]
```

Out[•]=

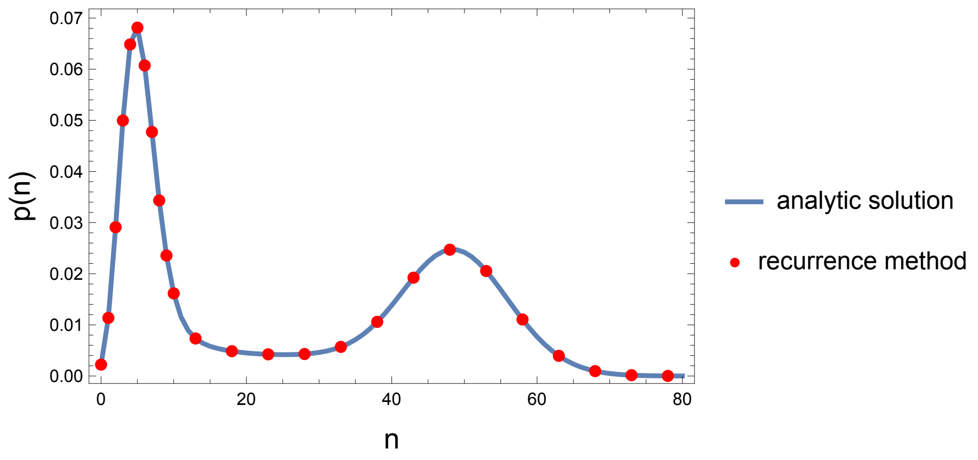


### Three-state model

#### Definitions

```
(*Implement recurrence relation given in Equations (28)-(33)*)
d[n_] := δ (δ^2 n^2 + δ n Sum[v_{i,j}, {i, 3}, {j, Drop[Range[3], {i}]}]
     + v_{1,2} (v_{2,3} + v_{3,1} + v_{3,2}) + v_{1,3} (v_{2,1} + v_{2,3} + v_{3,2}) + v_{2,1} (v_{3,1} + v_{3,2}) + v_{2,3} v_{3,1})
```

In[•]:=
```
(* initial conditions*)
nn = + v_{1,2} (v_{2,3} + v_{3,1} + v_{3,2}) + v_{1,3} (v_{2,1} + v_{2,3} + v_{3,2}) + v_{2,1} (v_{3,1} + v_{3,2}) + v_{2,3} v_{3,1};
ini = 1/nn Prepend[Table[Sum[v_{j,k} v_{k,i} /. j → Complement[Range[3], {i, k}][[1]],
        {k, Drop[Range[3], {i}]}] + v_{j,i} v_{k,i} /.
      {k → Drop[Range[3], {i}][[1]], j → Drop[Range[3], {i}][[2]]}, {i, 3}], 0];
```

```
In[•]:= (* recurrence relation *)
hSymList = {h0, h1, h2};
recurrence := Join[Table[ji = Complement[Range[3], {i}][[1]];
   ki = Complement[Range[3], {i}][[2]];
   hSymList[[i]][n] == 1/(n d[n])
     (kk_i (δ^2 n^2 + δ n (v_{ji,i} + v_{ji,ki} + v_{ki,i} + v_{ki,ji}) + v_{ki,ji} v_{ji,i} + v_{ji,ki} v_{ki,i} + v_{ji,i} v_{ki,i})
        hSymList[[i]][n - 1]
       + kk_ji (δ n v_{ji,i} + v_{ji,ki} v_{ki,i} + v_{ji,i} (v_{ki,ji} + v_{ki,i})) hSymList[[ji]][n - 1]
       + kk_ki (δ n v_{ki,i} + v_{ji,ki} v_{ki,i} + v_{ji,i} (v_{ki,ji} + v_{ki,i})) hSymList[[ki]][n - 1]),
   {i, 3}],
  Table[hSymList[[i]][0] == ini[[i + 1]], {i, 3}]]
```

```
In[•]:= (*construct pmf from h list according to Equation (27) *)
p[h_, n_] := 1/n! Sum[Pochhammer[i, n] h[[n + i]] (-1)^(i+1), {i, 1, Length[h] - n - 1}]
```

### Example - Plot Figure 7 (b)

```
In[•]:= (*set parameter values*)
δ = 1;
v_{1,2} = 45/1000;
v_{2,1} = 35/1000;
v_{3,1} = 15/1000;
v_{1,3} = 45/1000;
v_{2,3} = 15/1000;
v_{3,2} = 35/1000;
kk_1 = 5;
kk_2 = 30;
kk_3 = 60;
```

```
In[•]:= (*perform recurrence relation*)
nest = RecurrenceTable[recurrence, {h0, h1, h2}, {n, 0, 400}];

(*construct pmf*)
hList = Plus @@ Transpose@nest;
pmf = p[hList, #] & /@ Range[0, 100] // N;
```

In[•]:=
```
(*plot results*)
pl2 = ListLinePlot[pmf, PlotRange → All, PlotStyle → Thickness[0.008],
  Frame → True, PlotLegends → {"recurrence method"},
  FrameLabel → {Style["n", 15], Style["p(n)", 15]}]
```

Out[•]=

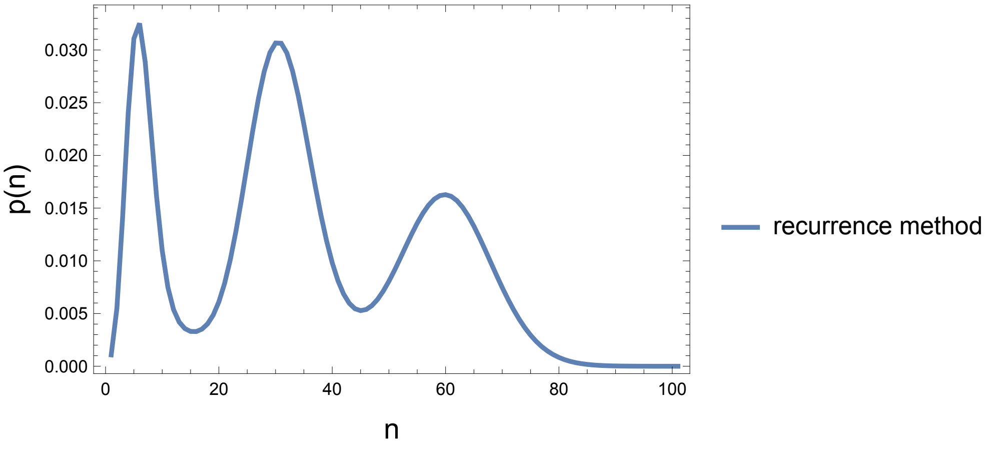


### Feedback model

#### Definitions

In[•]:=
```
(*construct pmf from h list according to Equation (27) *)
p[h_, n_] := 1/n! Sum[Pochhammer[i, n] h[[n + i]] (-1)^(i+1), {i, 1, Length[h] - n - 1}]
```

$$p[h\_, n\_] := \frac{1}{n!}\,\mathrm{Sum}\big[\mathrm{Pochhammer}[i, n]\; h[[n+i]]\; (-1)^{i+1},\ \{i, 1, \mathrm{Length}[h] - n - 1\}\big]$$

---

#### Parameters

In[•]:=
```
(*Definition of parameters as in Equations (A4)-(A5) and (A18)-(A20) *)
```

$$\Sigma_b := 1 + \sigma_b;$$

$$Rr := \rho_u - \rho_b\,\Sigma_b;$$

$$\alpha := \theta + \sigma_u \big/ Rr * (\rho_u - \rho_b);$$

$$\beta := 1 + \theta + 1 / \Sigma_b \left(\sigma_u + \rho_u \frac{\Sigma_b - 1}{\Sigma_b}\right);$$

$$w[z\_] := Rr\,\frac{\Sigma_b\, z - 1}{\Sigma_b{}^2};$$

### Analytic solution

In[•]:= (*analytic solution Equation (A23)*)

$$\text{pAna}[n\_] := \frac{\text{Aa}}{n!} \text{Sum}\Big[\text{Binomial}[n, m]\, \rho_b{}^{n-m}\, E^{-\rho_b} \frac{\text{Rr}^m\, \text{Pochhammer}[\alpha, m]}{\Sigma_b{}^m\, \text{Pochhammer}[\beta, m]} \left(\frac{\Sigma_b\, (\alpha + m)}{\sigma_b\, \rho_u} \text{Hypergeometric1F1}[\alpha + m + 1, \beta + m, w[0]] + \left(1 + \frac{\theta - \alpha}{\rho_u - \rho_b}\right) \text{Hypergeometric1F1}[\alpha + m, \beta + m, w[0]]\right), \{m, 0, n\}\Big]$$

### Initial conditions

In[•]:= (*Equation (A21)*)

$$\text{Aa} := \left(\frac{\Sigma_b}{\sigma_b} \frac{\alpha}{\rho_u} \text{Hypergeometric1F1}[\alpha + 1, \beta, w[1]] + \left(1 + \frac{\theta - \alpha}{\rho_u - \rho_b}\right) \text{Hypergeometric1F1}[\alpha, \beta, w[1]]\right)^{-1};$$

In[•]:= (*Initial conditions from Equations (A24) and (A25)*)

$$\text{h0Ini} := \text{Aa} \left(\frac{\Sigma_b}{\sigma_b} \frac{\alpha}{\rho_u} \text{Hypergeometric1F1}[\alpha + 1, \beta, w[1]] + \left(\frac{\theta - \alpha}{\rho_u - \rho_b}\right) \text{Hypergeometric1F1}[\alpha, \beta, w[1]]\right);$$

$$\text{h1Ini} := \text{Aa}\, \text{Hypergeometric1F1}[\alpha, \beta, w[1]];$$

$$\text{h01Ini} := \frac{1}{\sigma_b\, 1} (\theta + \sigma_u)\, \text{h0Ini};$$

### Round initial conditions

In[•]:= (*round to nn digit precision*)

$$\text{nn} = 800;$$

$$\text{h00} := \text{Round}\left[10^{\text{nn}}\, N[\text{h0Ini}, \text{nn}]\right] / 10^{\text{nn}};$$

$$\text{h10} := 1 - \text{h00};$$

In[•]:=

$$\text{h01} := \frac{1}{\sigma_b\, 1} \left((\theta + \sigma_u)\, \text{h10} + \sigma_u\, 0\, \text{h1}[n - 2] - (\sigma_b + 1)\, (1 - 1)\, \text{h0}[n - 1] + \rho_u\, 0\, \text{h0}[n - 2]\right);$$

### Recurrence equation

```
(*Equations (A14)-(A15)*)
eqs := {h0[n] == FullSimplify[
      1/(σb n) ((θ + σu) h1[n - 1] + σu h1[n - 2] - (σb + 1) (n - 1) h0[n - 1] + ρu h0[n - 2])],
    h1[n] == FullSimplify[(ρb + σu)/n h1[n - 1] - (σb + 1) h0[n] + ρu/n h0[n - 1]],
    h0[0] == h00, h1[0] == h10,
    h0[1] == h01};
```

### Example - Plot Figure 8

In[•]:=
```
(*set parameter values*)
ρu = 60;
ρb = 5;
θ = 0/100;
σu = 5/10;
σb = 4/1000;

(*perform recurrence relation*)
res = RecurrenceTable[eqs, {h0, h1}, {n, 0, 300}];

(*construct pmf*)
hList = (res[[ ;; , 1]] + res[[ ;; , 2]]);
resRecurrence = p[hList, #] & /@ Range[0, 90];

(*compute analytic solution*)
resAnalytic = pAna[#] & /@ Range[0, 90];
```

In[•]:=
```
(*plot results*)
pl1 = Show[
  ListLinePlot[resAnalytic, PlotRange → All, Frame → True,
   FrameLabel → {Style["n", 15], Style["p(n)", 15]}, PlotStyle →
    Thickness[0.008], Frame → True, PlotLegends → {"analytic solution"}],
  ListPlot[resRecurrence, PlotRange → All, PlotStyle → {Red, PointSize → 0.015},
   PlotRange → All, PlotLegends → PointLegend[{Red}, {"recurrence method"}]]]
```

Out[•]=

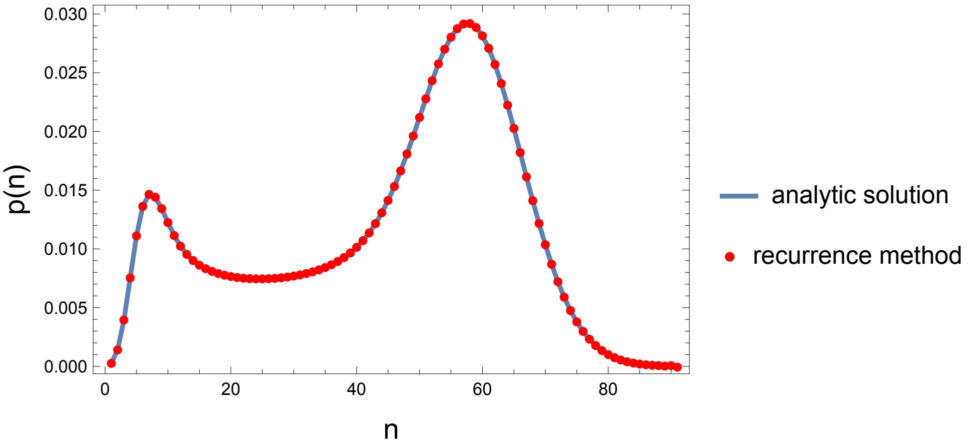
